# Improved optogenetic modification of the spiral ganglion neurons for future optical cochlear implants

**DOI:** 10.1101/2024.10.12.617982

**Authors:** Anupriya Thirumalai, Jana Henseler, Marzieh Enayati, Kathrin Kusch, Roland Hessler, Tobias Moser, Antoine Tarquin Huet

## Abstract

Optogenetic stimulation has become a promising approach for restoring lost body function. For example, partial restoration of vision has been achieved in a blind patient and proof-of-concept has been demonstrated for optogenetic hearing restoration in rodents. In order to prepare clinical translation of hearing restoration, efficient and safe optogenetic modification of spiral ganglion neurons (SGNs) in the mature cochlea remains to be developed. Here, we established microcatheter-based administration adeno-associated virus (AAV) to scala tympani of the cochlea of Mongolian gerbils and compared it to the previously developed AAV-injection into the spiral ganglion. We probed the potential AAV-PHP.S capsid to express channelrhodopsins (ChRs) under the control of the human synapsin promotor in mature SGNs in hearing and deafened gerbils. Using the microcatheter approach, but not with the AAV-modiolus injection, we achieved reliable ChR expression in SGN enabling optogenetic stimulation of the auditory pathway in 80% of the treated animals. Yet, the efficiency of SGN transduction was modest with only ∼30% ChR-expressing SGNs. Moreover, we encountered off-target expression in hair cells in hearing gerbils in both approaches, but not ChR expression in the central nervous system using microcatheter administration. Comparing optogenetic auditory brainstem responses of gerbils with and without hair cell transduction confirmed that SGNs were the primary site of optogenetic stimulation of the pathway.

## Introduction

More than 5% of the world’s population suffers from disabling hearing impairments, and the majority of them are affected from what is known as sensorineural hearing loss (WHO, 2021). In these patients, the hair cells that transduce the sound into micromechanical amplification (outer hair cells) or release of glutamate (inner hair cells, IHC) that activates the spiral ganglion neurons (SGN) are dysfunctional or absent. Despite recent clinical advances in cell therapy for otoferlin-related deafness (Qi *et al*, 2024; Lv *et al*, 2024), hearing aids and cochlear implants will remain the pillar of hearing restoration for the foreseeable future because of their one-size-fits-all nature. Hearing restoration for severely hearing-impaired patients relies on the use of electrical cochlear implants (eCI), that bypasses the lost or dysfunctional cochlear hair cells and directly stimulates the SGNs that are typically maintained for longer. By restoring hearing perception to the vast majority of its currently more than a million users, the eCI is considered to be the most successful neuroprosthesis to date (for review, see (Lenarz, 2018)). While eCI enables the perception of speech in a quiet environment, users recognize an unmet medical need in the real-life condition where background noise competes with speech (Hunniford *et al*, 2023; Wolf *et al*, 2022). This deficiency is commonly attributed to the wide current spread from each electrode contact in the saline-filled cochlea, activating large SGN groups and limiting spectral information transfer (Kral *et al*, 1998). Despite engineering efforts to reduce the spread of electrical stimulation, clinical benefits have remained limited. An alternative way to achieve a spatially narrower activation of the SGNs, and thus to increase the spectral information transfer, is to switch the stimulation modality to light, since light can be better spatially confined (Dieter *et al*, 2019, 2020; Keppeler *et al*, 2020). In this approach, photocontrol is achieved by driving viral vector-mediated expression of channelrhodopsin (ChR) in the SGNs. This concept leads to the current development of an optical cochlear implant (oCI) consisting of a medical device on the one hand and an optogene therapy on the other (for review, see (Huet *et al*, 2024)).

For the oCI to be translated into a clinical approach, efficiency, reliability and safety of optogene therapy needs to be demonstrated in preclinical work. Previous work has shown that efficient optogene therapy can be achieved by injection of a viral vector carrying ChR under the control of the human synapsin promoter (hSyn) into *scala tympani* of early postnatal rodents (i.e. “intrascalar”, (Mager *et al*, 2018; Keppeler *et al*, 2018; Huet *et al*, 2021; Richardson *et al*, 2021; Duarte *et al*, 2018). At this age of optogene therapy, a broad range of viral capsids was evaluated and reached similar rates of ChR expression in the SGNs: AAV2/6 (Mager *et al*, 2018; Bali *et al*, 2022; Huet *et al*, 2021; Wrobel *et al*, 2018), AAV2/9 (Zerche *et al*, 2023), AAV PHP.B (Keppeler *et al*, 2018; Huet *et al*, 2021; Bali *et al*, 2021; Keppeler *et al*, 2020; Mittring *et al*, 2023; Dieter *et al*, 2019, 2020; Michael *et al*, 2023), AAV PHP.eB (Bali *et al*, 2021; Michael *et al*, 2023) and AAV.Anc80 (Duarte *et al*, 2018; Richardson *et al*, 2021). More recently, early postnatal optogene therapy was demonstrated to be stable for two years (the lifetime of mice) but associated with extracochlear ChR expression at different levels of the central nervous (Bali *et al*, 2022). ChR expression in SGN adult cochleae can also be achieved by direct injection of the viral vector into the cochlear modiolus (“intramodiolar”, directly into the Rosenthal canal of the bony modiolus that houses the spiral ganglion, (Wrobel *et al*, 2018)). The success of the optogene therapy for enabling optogenetic SGN stimulation at this age was limited to ∼40% of the treated gerbils and a low ChR expression rate (∼20-30% SGNs) in the responsive cochleae (Wrobel *et al*, 2018; Huet *et al*, 2021)). In addition, intramodiolar pressure injection caused a loss of ∼20% of the SGNs (Wrobel *et al*, 2018). Even less efficient and reliable transduction was reported upon intrascalar AAV injection in adult animals (Richardson et al, 2021).

In this study, we aimed to develop a translatable delivery approach and to identify appropriate viral vectors that would allow ChR expression in the SGNs of all treated adult animals as relevant for future clinical work. Previous work in adult mice demonstrated the role of the cochlear aqueduct in the spread of drugs locally administered in the cochlea towards the brain (Talaei *et al*, 2019). For this purpose, we selected the capsid AAV.PHP-S for its tropism towards the peripheral nervous system (Chan *et al*, 2017), thus avoiding the transduction of neurons in the central nervous system in the event that the viral suspension would spread towards the brain. First, we evaluated the capsid AAV.PHP-S for its ability to mediate ChR expression in young and adult SGNs employing the Mongolian gerbil for its large cochlea (Keppeler *et al*, 2021). We then tested multiple administration approaches that allow for the replacement of cochlear fluids in a standardized and controllable manner and compared them to the currently established optogene therapy for adult animals via intramodiolar injection. We found that slow delivery of AAV.PHP-S via a microcatheter inserted into the round window in combination with an evacuation vent at the oval window (RW_µ-cat_ + vent) allowed expression of ChR in all treated cochleae and that optically evoked auditory potentials could be recorded in >80% of the treated cochleae. Finally, in a pharmacological model of profound sensorineural hearing loss, we demonstrated that successful optogene therapy by AAV-mediated transduction of SGNs by RW_µ-cat_ + vent administration does not require the presence of IHCs.

## Results

### AAV.PHP-S as a viral vector for optogenetic modification of spiral ganglion neurons

Here, we evaluated AAV.PHP-S, a capsid designed to specifically transduce neurons from the peripheral nervous system (Chan et al, 2017), for its ability to mediate ChR expression in the SGNs. AAV.PHP-S was first evaluated following pressure injection into the cochlea of early postnatal age gerbils (**figure 1.B**), which, at this age, is associated with high SGN transduction rate in most of injected gerbils (Huet *et al*, 2021). We expressed the blue-light activated ChR2-variant, CatCh (Kleinlogel *et al*, 2011; Wrobel *et al*, 2018), fused to the enhanced yellow fluorescent protein (eYFP) under control of hSyn promoter (**figure 1.A**). The capsid was injected at a titer of 6.42 x 10^12^ genome copies per milliliter (gc/mL). Approximately, 8 weeks after injection, expression of CatCh-eYFP was analysed by confocal microscopy of cross-modiolar section immunolabeled for GFP, parvalbumin as a SGN marker and calretinin as an inner hair cell (IHC) marker, following functional characterization regardless of the presence or absence of optically evoked optically evoked auditory brainstem responses (oABRs, **figure 1.C-D**) that reflect the optical activation of the peripherical auditory pathway, and for, both, the injected and non-injected cochlea. To observe SGNs and the IHCs they innervate, we developed the semi-thick cross-modiolar section of the cochlea (thickness = 220 µm, **figure 1.A**). For efficient analysis of the large number of cells in the images, the SGNs were automatically segmented in 3 dimensions using a self-trained model in CellPose 2.0 (Pachitariu & Stringer, 2022) of Arivis 4D software (see the section “Material and methods” for details). ChR-expressing SGNs (GFP^+^ SGNs) were semi-automatically identified based on the distribution of averaged GFP signal per 3D segmented cell (Huet *et al*, 2021; Mittring *et al*, 2023).

**Figure 1.**
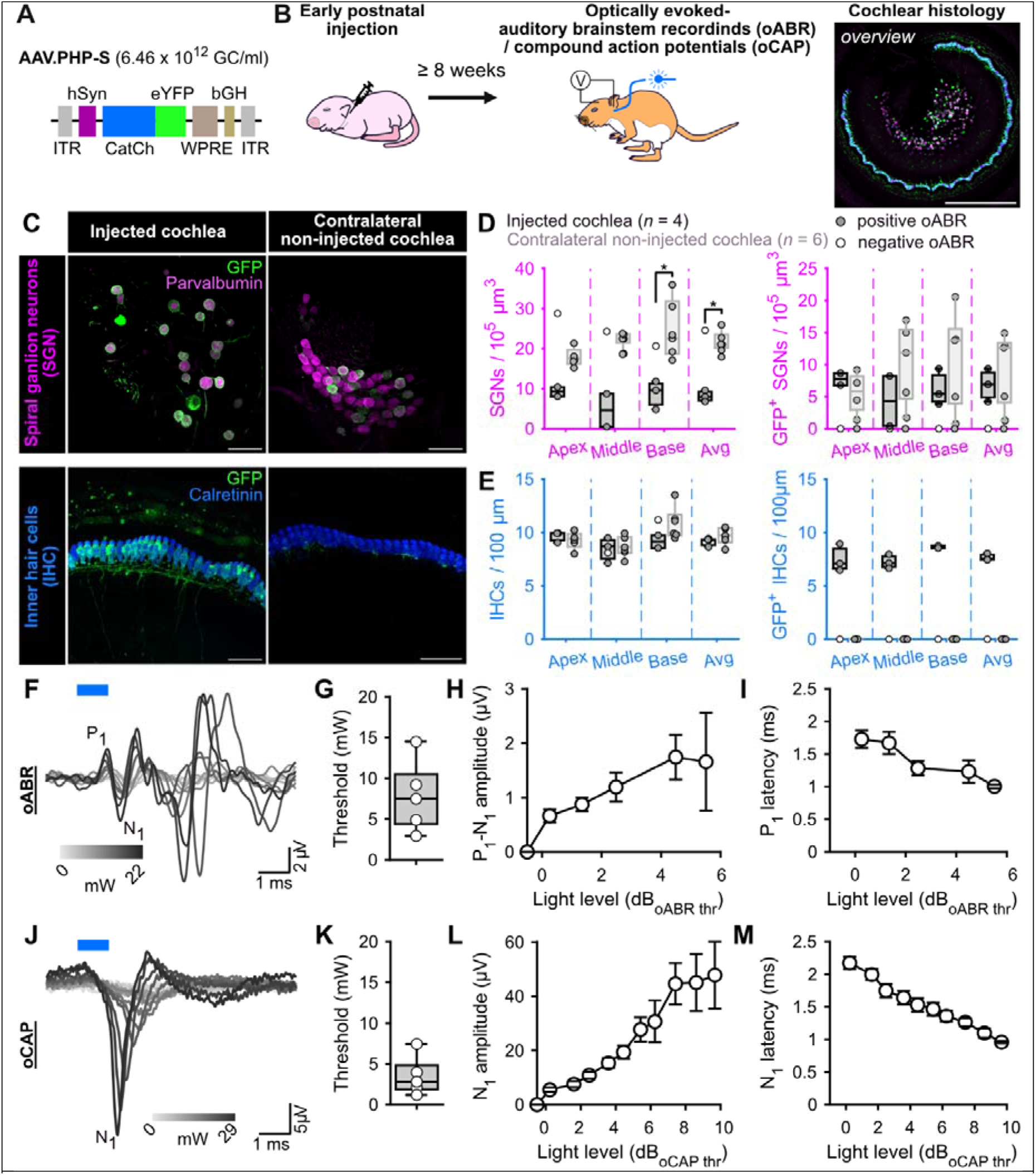
AAV.PHP-S mediates efficient ChR-expression of CatCh in SGNs following cochlear injection at early postnatal age. **A.** pAAV plasmid used in this study containing CatCh fused with eYFP. Expression was driven by human synapsin promoter (hSyn) and enhanced by the Woodchuck hepatitis virus post-translational regulatory element (WPRE) and bovine growth hormone (bGH). ITR corresponds to the inverted terminal repeats. **B.** Time course of the experiment: At early postnatal age, animals received an intracochlear viral suspension injection. At least 8 weeks later, oABRs were measured and the cochleae were collected for histology. **C.** Representative maximum projection of confocal images obtained from immunolabelled cross-modiolar sections of an injected (left) and a non-injected cochlea (right, scale bar = 50 µm). GFP (green) marks ChR-expressing cells. The first raw corresponds to the spiral ganglion neurons (SGN) labelled with parvalbumin (magenta). The second raw corresponds to the inner hair cells (IHC) labelled with calretinin (blue). **D-E.** Quantification of the SGN density, GFP^+^ SGN density (D), IHC density and GFP^+^ IHC density (E). Data from the injected cochleae are represented in black, and contralateral non-injected cochlea in grey. For the injected cochlea, a grey-filled maker was used when positive oABRs were measured and an open-marker for the negative oABRs. Wilcoxon rank sum test (*, *P* ≤ 0.05). **F.J.** Representative optically evoked auditory brainstem recordings (oABR, F) and optically evoked compound action potentials (oCAP). The first wave of both potentials reflects the synchronous activation of the SGNs. **G-I, K-M.** Quantification of the activation thresholds (G,K), first wave amplitude (H,L) and first wave latency (I,M) of the oABR (G-I, *n* = 5 cochleae) and the oCAP (K-M, *n* = 5 cochleae). The potential latencies and amplitudes are expressed as mean ± SEM as a function of the light level above threshold (see Material and methods for details). Box plots show minimum, 25^th^ percentile, median, 75^th^ percentile, and maximum.

In 5 out of the 6 injected cochleae, oABRs could be elicited and ChR-expressing SGNs were found. In the negative cochlea, no ChR-expressing SGNs were found and thus, we hypothesized that this cochlea must have been mis-injected and its data will only be displayed but not included in the following quantification. The SGN density of the injected cochleae was significantly reduced compared to the contralateral non-injected cochleae (**figure 1.C-D**, injected: 8.12 ± 0.81, *n* = 3 and non-injected 21.73 ± 1.31 SGNs / 10^5^ µm^3^, *n* = 5, *P* = 0.036, Wilcoxon rank). Our previous study reported a slight decrease of SGNs, potentially induced by a transient increase of pressure into the cochlea during the injection (Huet *et al*, 2021), but the reduction was stronger in the current study.

The density of GFP^+^ SGNs was similar between the transduced cochleae and amounted to 6.83 ± 1.49 and 9.35 ± 2.63 SGNs / 10^5^ µm^3^ for injected and non-injected cochlea, respectively (**figure 1.C,D**). To take into account the difference of SGNs density between both sides, ChR-expression was also quantified as the ratio between the number of GFP^+^ SGNs and the total number of residual SGNs (i.e., the ChR-expression rate). In this case, the transduction rate tended to be higher for the injected cochleae (82.17% ± 10.54) compared to the non-injected ones (45.3% ± 13.08, *P* = 0.14, Wilcoxon rank sum test).

Contrary to previous reports, IHCs were also expressing CatCh-eYFP and thus we quantified the extent of their ChR-expression. The IHC density was similar between both ears and amounted to 9.18 ± 0.18 and 9.64 ± 0.21 IHC / 100 µm for the injected (*n* = 5) and the contralateral non-injected (*n* = 6) cochleae. Nonetheless, ChR-expressing IHCs were solely limited to the injected cochlea (7.67 ± 0.15 GFP^+^ IHC / 100 µm, i.e. ChR-expression rate = 85.47 ± 1.47%). We assume that the cross-modiolar slicing facilitates the detection of ChR-expressing IHCs, compared to the mid-modiolar sections used in the previous studies.

Next, we recorded oABRs from scalp electrodes in response to 1 ms light pulses delivered into the cochlea, at 10 Hz, by an optical fiber (Iii = 200 µm) coupled to a 488 nm laser (**figure 1.B**). At low light intensities, the oABRs were characterized by 2-3 positive waves as previously described in gerbils (Wrobel *et al*, 2018; Dieter *et al*, 2019, 2020; Huet *et al*, 2021). At higher light intensities, the oABRs were contaminated by a large and late likely non-auditory potential, potentially reflecting facial nerve activation (**figure 1.F**). The oABR threshold on average amounted to 7.8 ± 1.99 mW (**figure 1.G**). To take into account the large distribution of oABR thresholds, quantification of the oABR wave amplitudes and latencies was related to light levels relative to the oABR threshold using the following formula: Light level (dB_oABR threshold_) = 10 x log_lO_(A⁄A_threshold_) where A is the radiant flux and A_threshold_ the radiant flux at the oABR threshold in mW. The amplitude of the first oABR wave increased linearly with the light level (**figure 1.H**) while the first wave latency decreased (**figure 1.I**) up to ∼4 dB. To characterize SGN responses to a wider range of light levels regardless of the absence or presence of non-auditory potentials, optically evoked compound action potentials (oCAP) were recorded using an electrode placed at the surface of the cochlea (**figure 1.J**). The oCAPs reflect the synchronous activation of the SGNs and are recorded from a silver ball electrode implanted in the RW niche (Bourien *et al*, 2014; Batrel *et al*, 2017). The oCAP thresholds tended to be lower than those of oABRs measured from the same animal and amounted to 3.5 ± 1.09 mW (**figure 1.K**, *P* = 0.06, *n* = 5, Wilcoxon signed rank test). The extent of cochlear activation, reflected by the oCAP wave amplitude, increased with the radiant flux for up to 7 dB and seemed to saturate above (**figure 1.L**). Nonetheless, the oCAP latency decreased over the whole tested range (> 9 dB, **figure 1.M**).

The SGN loss combined with the IHC-transduction in the injected cochlea suggested that the titer of AAV-PHP.S-hSyn-CatCh-eYFP was too high for achieving safe and specific optogene therapy when administered at early postnatal age. The off-target expression in IHCs might not be clinically relevant given loss of hair cells in sensorineural hearing loss. Nonetheless, the data indicate that promotor choice - synapsin is not expressed at the protein level in IHCs (Safieddine & Wenthold, 1999) – does not ensure specificity provided sufficient AAV titer. Future studies will be needed to identify the optimal AAV titer for efficiently, specifically and safely expressing ChRs in SGNs at that age.

### Development of RW_µ-cat_ + OW approach to improve the reliability of optogene therapy in adults

While AAV injection into the *scala tympani* at early postnatal age results in reliable transduction and ChR expression of SGNs, adult intramodiolar injection is characterized by a poorer outcome, with SGN transduction being achieved in only a fraction of the treated cochlea (Huet *et al*, 2021; Wrobel *et al*, 2018; Dieter *et al*, 2019; Michael *et al*, 2023). Moreover, while surgically feasible in humans (Wrobel *et al*, 2021), a round window approach might represent the first choice, given its frequent use in CI surgery, its successful use for clinical cochlear gene therapy targeting IHCs (Lv *et al*, 2024; Wang *et al*, 2024) and the availability of approved microcatheters (µ_cat_). However, to our knowledge a systematic preclinical SGN gene therapy study had yet to be performed. We hypothesized that more reliable SGN transduction could be achieved by administering AAV to the *scala tympani* that has a ∼18x larger volume (1.81 µL in the gerbil) than Rosenthal’s canal (0.10 µL) that largely occupied by SGNs (Keppeler *et al*, 2021).

Here, we tested four distinct approaches employing micro-pump-driven AAV administration via a µ_cat_ provided by MED-EL (figure 2**.A**). The following approaches were tested: *i*) microcatheter (µ-cat) insertion via the round window (RW_µ-cat_); *ii*) µ-cat and pressure relief by a vent at the oval window (RW_µ-cat_ + OW); *iii)* µ-cat _t_ and pressure relief in the posterior semi-circular canal (RW_µ-cat_ + PSCC); *iv*) PSCC delivery and pressure relief evacuation vent drilled at the RW (PSCC + vent). In a preliminary set of experiments, those different approaches (RW_µ-cat_, *n* = 5; RW_µ-cat_ + PSCC, *n* = 7; RW_µ-cat_ + OW, *n* = 5; PSCC_cat_ + vent, *n* = 6) were compared to the reference direct pressure modiolus injection (*n* = 6, **figure S1.A**) using AAV2/9-hSyn-f-Chrimson-eYFP (titer = 1-3 x 10^12^ GC/mL). In all those approaches, the viral suspension was mixed with fast-green (1:20) to visualize the evacuation of the suspension. AAV-administration was performed at 250 - 300 nL/min using a micropump and was stopped either after observing reflux of the suspension or after dosing 5 µL.

**Figure 2.**
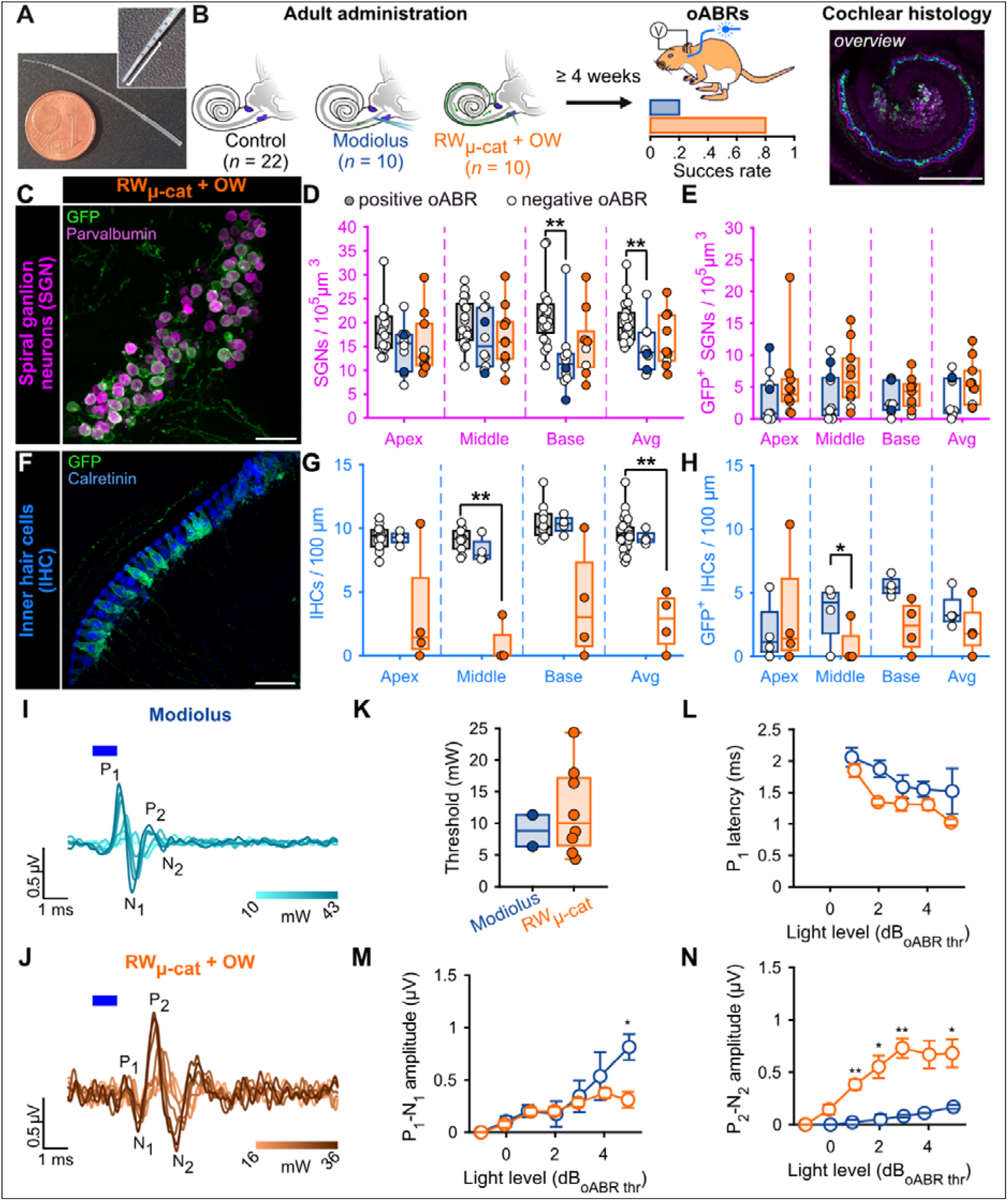
Viral administration with a micro-catheter inserted at the round window + vent at the oval window (RW_µ-cat_ + OW) mediates improved optogenetic modification of the SGNs compared to the reference modiolus injection. **A.** Picture of the catheter used for the RW_µ-cat_ + OW approach. The white line indicates the insertion depth (4.5-5 mm). **B.** Time course of the experiment. At adult age, animals were divided in three groups: i) control: no injection (black); ii) modiolus injection (blue); iii) RW_µ-cat_ + OW administration (orange) of AAV-PHP.S-hSyn-CatCh. At least 4 weeks later, oABRs were measured. The proportion of animals for which a positive oABR were recorded are represented as boxplot. Next, the cochleae were collected for histology. Here, the overview image is a cross-modiolar section from a modiolus injected cochlea (scale bar = 500 µm). **C,F.** Representative maximum projection of confocal images obtained from immunolabelled cross-modiolar sections of spiral ganglion neurons (SGN, C, scale bar = 50 µm) and inner hair cells (IHC, F, scale bar = 50 µm). GFP (green) marks ChR-expressing cells. The SGNs were labelled with parvalbumin (magenta). The IHCs were labelled with calretinin (blue). **D-E, G-H.** Quantification of the SGN density (D, *n* = 6), GFP+ SGN density (E, *n* = 6), IHC density (G, *n* = 4) and GFP+ IHC density (H, *n* = 4). A filled marker was used when positive oABRs were measured and an open-marker for the negative oABRs. Kruskal-Wallis test followed by a multi-comparison test (*, *P* ≤ 0.05; **, *P* ≤ 0.01). **I-J.** Representative oABRs recorded from modiolus injected (I) and RW_µ-cat_ + OW administered cochleae. The light intensity is color coded using the color scale in insert. **K.** Quantification of the oABR activation threshold measured from modiolus injected (orange, *n* = 2 positive oABR cochleae out of 10 injected ones) and RW_µ-cat_ + OW administred (blue, *n* = 8 positive oABR cochleae out of 10 injected ones) cochleae. **L-N.** Quantification of the P_1_ latency (L), P_1_-N_1_ amplitude (M) and P_2_-N_2_ amplitude (N) as a function of the light level relative to the oABR threshold. Box plots show minimum, 25^th^ percentile, median, 75^th^ percentile, and maximum. Averaged ± SEM. Wilcoxon rank sum test (*, *P* ≤ 0.05; **, *P* ≤ 0.01).

Approximately, 4 weeks after injection, animals were tested and expression of f-Chrimson-eYFP was analyzed by confocal microscopy of mid-modiolar cryosection immunolabeled for GFP and parvalbumin as a SGN marker, regardless of the presence or absence of optically evoked oABRs (**figure S1.B-C**). RWµ-cat + vent administrations tended to enable higher SGN and GFP^+^ SGN densities and expression rates compared to the reference modiolus injection (table 1, **figure S1.B**) (table 1, **figure S1.C**).

**Table 1.**

| | SGN density<br>(SGN/$10^5 \mu\text{m}^3$) | GFP<sup>+</sup> SGN density<br>(SGN/$10^5 \mu\text{m}^3$) | Absolute expression<br>rate (%) |
| --- | --- | --- | --- |
| Modiolus | 11.61 ± 0.78 | 1.04 ± 0.79 | 8.05 ± 5.66 |
| RW<sub>μ-cat</sub> | 16.85 ± 1.82 | 0.72 ± 0.49 | 2.86 ± 0.89 |
| RW<sub>μ-cat</sub> + PSSC | 16.59 ± 0.97 | 2.11 ± 0.65 | 12.79 ± 3.21 |
| RW<sub>μ-cat</sub> + OW | 13.69 ± 1.52 | 1.52 ± 0.34 | 11.90 ± 2.60 |
| PSSC<sub>cat</sub> + vent | 19.17 ± 1.46 | 1.14 ± 0.5 | 1.53 ± 1.02 |

In order to further scrutinize the efficacy of the RW_µ-cat_ + OW delivery approach over the modiolus injection for optogenetic modification of the SGNs, we compared both approaches on a larger sample of animals (*n* = 10 for both groups) using AAV-PHP.S-hSyn-CatCh-eYFP (titer = 6.4 x 10^12^ GC/mL) against control non-injected cochleae (*n* = 22). Approximately, 4 weeks after administration, we tested for oABRs and expression of CatCh-eYFP was analysed by confocal microscopy of either mid-modiolar cryosection (*n* = 6 for both groups) or cross-modiolar section (*n* = 4 for both groups) immunolabeled for GFP, parvalbumin and calretinin (only for the cross-modiolar sections), regardless of the presence or absence of optically evoked oABRs (figure 2**.B**). The SGN density was reduced for both approaches compared to the control, figure 2.C-D, table 2) and was significantly lower for the modiolus injected group (*P* = 0.0093, Kruskal-Wallis test followed by a Tukey-Kramer post-hoc test). The number of SGN expressing CatCh-eYFP was almost twice bigger following RW_µ-cat_ + vent administration compared to the modiolus injected cochleae (figure 2.C,E, table 2). Interestingly, a relative transduction rate of at least 20% was found in only 40% of the modiolus injected cochleae compared to 70% of the cochleae treated via RW_µ-cat_ + vent.

**Table 2.**
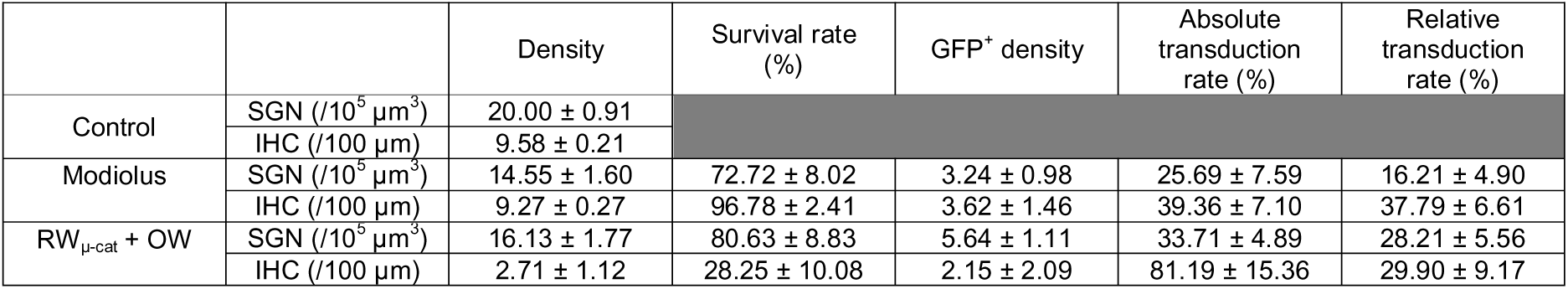

Here again, IHCs expressing CatCh-eYFP were found in both injection groups and thus IHC histology was performed from cochlear cross-modiolar slices. Quantification of the IHC density revealed a drastic loss of hair cells for the cochleae that underwent a RW_µ-cat_ + OW administration compared to the control cochleae (**table 2**, *P* = 0.0027, Kruskal-Wallis test followed by a multi-comparison test). This most likely reflects the insertion trauma caused by the micro-catheter insertion into the *scala tympani*. oABRs were found in 20% of the animals subjected to intramodiolar injection and and 80% of the animals undergoing RW_µ-cat_ + OW administration. oABRs activation thresholds were similar between both groups (8.85 ± 2.49 and 12.00 ± 2.46 mW for modiolus and RW_µ-cat_ + vent, respectively) but oABRs strongly differed in term of wave morphologies. At 5 dB above the threshold, modiolus injected cochleae were characterized by a larger first oABR wave (P_1_-N_1_) than RW_µ-cat_ + vent administered cochleae (figure 2**.M**, 0.81 ± 0.12 and 0.31 ± 0.08 µV, *P* = 0.033, Wilcoxon rank sum test), but were also characterized by a ∼1 order of magnitude smaller second wave (P_2_-N_2_, figure 2**.M**, 0.17 ± 0.02 and 0.68 ± 0.13 µV, *P* = 0.017, Wilcoxon rank sum test).

The increased reliability of SGN transduction −80% of treated cochleae were oABR-positive – together with a stronger activation of the auditory pathway - the larger 2^nd^ oABR wave reflecting activation of the auditory brainstem - make of the RW_µ-cat_ + OW approach a stronger candidate route of administration for optogenetic modification of the SGNs than the previously established intramodiolar AAV injection.

### Validation of RW_µ-cat_ + OW administration to mediate the transduction of the SGN in adult deafened cochleae

The main indication for cochlear implantation is to restore hearing in patients with profound sensorineural hearing loss characterized by absent or dysfunctional IHCs. Therefore, we investigated whether the RW_µ-cat_ + vent administration could transduce SGN in absence of IHC. Induction of the profound hearing loss was achieved by injecting 3 µL of Kanamycin (100 mg/ml) into the cochlea by the RW one week before to RW_µ-cat_ + vent administration of AAV-PHP.S-hSyn-CatCh-eYFP (titer = 6.4 x 10^12^ GC/mL, *n* = 6 animals, figure 3**.A**).

**Figure 3.**
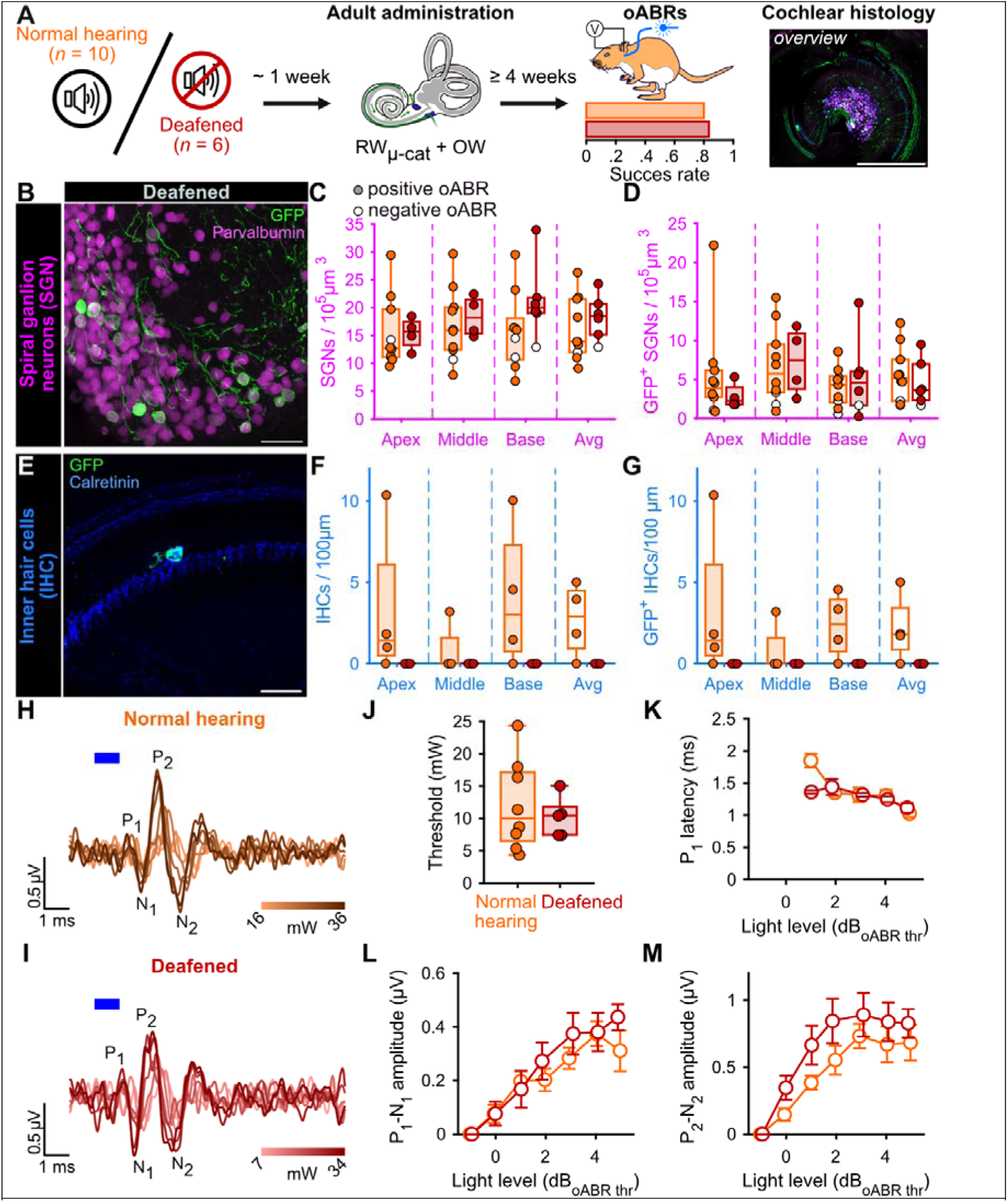
Viral administration with a RW_µ-cat_ + OW does not require the presence of inner hair cell to optogenetically modify the SGNs. **A.** Time course of the experiment. At adult age, a group of animals was deafened by cochlear round window injection of Kanamycin (100 mg/mL, *n* = 6). The control normal hearing animals are replotted from the figure 2. One week after deafening, animals received a RW_µ-cat_ + OW administration of AAV-PHP.S-hSyn-CatCh. At least 4 weeks later, oABRs were measured. The proportion of animals for which a positive oABR were recorded are represented as boxplot. Next, the cochleae were collected for histology. Here, the overview image is a cross-modiolar section from a deafened cochlea (scale bar = 500 µm). **B,E.** Representative maximum projection of confocal images obtained from immunolabelled cross-modiolar sections of spiral ganglion neurons (SGN, B, scale bar = 50 µm) and inner hair cells (IHC, E, scale bar = 50 µm). GFP (green) marks ChR-expressing cells. The SGNs were labelled with parvalbumin (magenta). The IHCs were labelled with calretinin (blue). **C-D, F-G.** Quantification of the SGN density (C), GFP+ SGN density (D), IHC density (F) and GFP+ IHC density (G). A filled maker was used when positive oABRs were measured and an open-marker for the negative oABRs. **H-I.** Representative oABRs recorded in normal hearing (H) and deafened (I) cochleae following RW_µ-cat_ + OW administration. The light intensity is color coded using the color scale in insert. **J.** Quantification of the oABR activation threshold measured from normal hearing (orange) and deafened (red) cochleae. **K-M.** Quantification of the P_1_ latency (K), P_1_-N_1_ amplitude (L) and P_2_-N_2_ amplitude (M) as a function of the light level relative to the oABR threshold. Box plots show minimum, 25^th^ percentile, median, 75^th^ percentile, and maximum. Averaged ± SEM.

Treated deafened cochleae were compared to the treated normal hearing cochleae described above (figure 2). The success of the deafening procedure was confirmed by the absence of IHC observed at any turn of the kanamycin-injected cochleae by confocal imaging following immunolabeling (figure 3**.E-G**). The extent of the optogenetic modification was identical between normal hearing and deafened cochleae as shown by comparable GFP^+^ SGN densities (**Table 2**, figure 3**.B-D**) and oABRs (figure 3**.H-M**). Moreover, the deafening protocol did not reduce the total SGN density. Together those data confirm the utility of RW_µ-cat_ + vent viral vector delivery to optogenetically modify the SGNs in the naive and deafened cochlea.

|  |  | Density | Survival rate (%) | GFP<sup>+</sup> density | Absolute transduction rate (%) | Relative transduction rate (%) |
| --- | --- | --- | --- | --- | --- | --- |
| Normal hearing RW<sub>μ-cat</sub> + vent | SGN (/10<sup>5</sup> μm<sup>3</sup>) | 16.13 ± 1.77 | 80.63 ± 8.83 | 5.64 ± 1.11 | 33.71 ± 4.89 | 28.21 ± 5.56 |
|  | IHC (/100 μm) | 2.71 ± 1.12 | 28.25 ± 10.08 | 2.15 ± 2.09 | 81.19 ± 15.36 | 29.90 ± 9.17 |
| Deafened RW<sub>μ-cat</sub> + vent | SGN (/10<sup>5</sup> μm<sup>3</sup>) | 18.33 ± 1.64 | 91.64 ± 8.21 | 4.63 ± 1.22 | 24.00 ± 4.78 | 23.16 ± 6.13 |
|  | IHC (/100 μm) | 0 ± 0 | 0 ± 0 | 0 ± 0 | 0 ± 0 | 0 ± 0 |

Finally, we examined a subset of 5 brains (3 normal hearing RWµ-cat + OW and 2 deafened RW_µ-cat_ + OW) for the off-target expression of ChR (**Figure S2**). Histological analysis was performed on coronal slices at the level where axons from SGNs project to cochlear nucleus neurons (i.e., the first brain region exposed to the viral suspension after cochlear administration, (Talaei *et al*, 2019)). Expression of GFP was restricted to the cochlear nucleus on the injected side and was confined to the large axosomatic synapses on bushy cells (i.e., endbulbs of Held). Although this observation does not substitute for more extensive histologic analysis of larger brain regions, it does support the absence of major spread of AAV and off-target expression of ChR transduction upon the RW_µ-cat_ + OW administration of AAVs.

### Functional impact of CatCh expression in both IHCs and SGNs

In previous work with administration AAV2/6, AAV-PHP.B and AAV-PHP.eB carrying ChRs under the control of hSyn promoter (titer between 2 x 10^12^ and 5 x 10^13^ GC/mL), the expression of ChR was limited to SGNs and no expression of ChR was reported in the IHCs (Mager *et al*, 2018; Keppeler *et al*, 2018; Huet *et al*, 2021; Bali *et al*, 2021; Wrobel *et al*, 2018). Here, using AAV-PHP.S with hSyn promoter at a similar titer (6.4 x 10^12^ gc/mL), we observed co-transduction of ChR in the IHCs in the majority of the treated adult cochleae. We therefore studied the opto-physiological relevance of IHC co-activation taking advantage of the comparison of cochlea with or without ChR-expressing IHCs.

Firstly, we identified that at least 2.34 SGN/10^5^ µm (11.7% of the total amount of SGN in non-treated cochleae) were required to elicit an oABR (figure 4**.A**). The mean number of ChR-expressing SGNs was similar between positive oABR cochleae with or without ChR-expressing IHCs: 8.47 ± 1.97 SGN/10^5^ µm and 6.05 ± 1.37, respectively. The number of ChR-expressing IHC did not differ between oABR-positive and - negative cochleae (figure 4**.B)**: 2.86 ± 1.86 and 2.26 ± 0.69 GFP^+^ IHC/100 µm, respectively. This suggests that oABRs are mainly caused by the photoactivation of the SGNs rather than IHCs.

**Figure 4.**
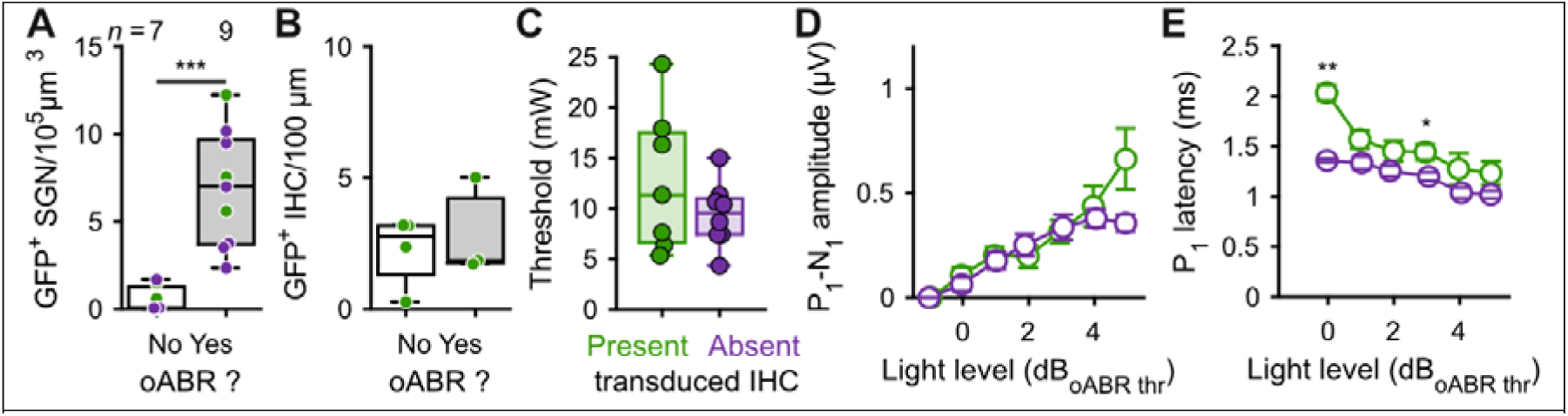
Contribution of optogenetically modified inner hair cell response to the oABRs. **A.** Quantification of the GFP^+^ SGN density as a function of the presence of the oABR response. **B.** For the cochleae where ChR-expressing IHC were observed, quantification of the IHC transduction as a function of the presence of the oABR. **C.** Quantification of the oABR threshold for cochlea where GFP+ IHC were present (green) or absent (purple). **D-E.** Quantification of the P_1_ latency (C), P_1_-N_1_ amplitude (D) as a function of the light level relative to the oABR threshold. Box plots show minimum, 25^th^ percentile, median, 75^th^ percentile, and maximum. Averaged ± SEM. Wilcoxon rank sum test (*, *P* ≤ 0.05; **, *P* ≤ 0.01).

Next, we investigated if the presence of ChR-expressing IHCs influenced the oABR properties. The oABR threshold radiant flux was similar for cochleae with and without ChR-expressing IHCs were observed (figure 4**.C**): 12.76 ± 2.66 mW and 9.42 ± 1.13, respectively. At all light levels, the oABR amplitudes were similar between the cochleae with and without ChR-expressing IHCs (figure 4**.D**). Nonetheless, the oABR first wave latency tended to be longer for the cochlea with ChR-expressing IHCs and this effect was the strongest near threshold (figure 4**.E**, 1.36 ± 0.02 and 2.03 ± 0.08, *P* = 0.009, Wilcoxon rank sum test).

## Discussion

Here we used mature Mongolian gerbils as a preclinical rodent model to further develop optogene therapy for the future optical cochlear implant. The Mongolian gerbil is a small rodent (∼ 60-80 g) with a hearing frequency range similar to that of humans (Huet *et al*, 2016). Its low-frequency hearing is supported by a relatively large middle ear and cochlea, which allows cochlear implant and drug delivery studies to be conducted in better resemblance to the human ear than if conducted in high-frequency hearing rodents (e.g., mice and rats). We identified the AAV administration via a round window-inserted microcatheter with a pressure relief via an oval window vent to provide a reliable means for optogene therapy. Roughly 80% of the treated animals showed optogenetic activation of the auditory pathway compared to 40% with the previously described intramodiolar AAV injection (this study and (Wrobel *et al*, 2018)). We identified substantial off-target expression in IHCs but no obvious ChR expression in the brain. Optogenetic stimulation of the auditory pathway required ≥10% of SGNs to express ChR and was robust to the absence of ChR expressing IHCs (e.g., lack of transduction or kanamycin deafening). Together this study advances optogenetic hearing restoration by demonstrating the utility of a clinically established AAV administration route and studying the impact that hair cell stimulation could have on optogenetic hearing restoration in cases with residual low frequency hearing.

### Old and new standards for evaluating SGN gene therapies

The standard routine to assess the activation of the SGNs *in vivo* consists to measure the auditory brainstem response (ABR). Acoustic and optical ABRs are characterized by 1 to 3 waves with the first wave reflecting the synchronous activation of the SGNs (Bourien *et al*, 2014; Melcher *et al*, 1996). Previous studies have shown that oABRs are a good proxy for ChR transduction of SGNs (Wrobel *et al*, 2018; Huet *et al*, 2021).

In previous studies developing optogenetic modification of SGNs, IHCs transduction was not observed in mid-modiolar cryosections of the cochlea, where only a few IHCs are seen per turn (Wrobel *et al*, 2018; Huet *et al*, 2021). Using AAV.PHP-S and scala tympani delivery, our initial observations suggested that transduction of IHCs was systematic (further discussed below). Therefore, we developed the semi-tick (220 µm) cross-modiolar sections to visualize the SGNs and the IHCs they innervate from the same section. This approach appeared to be a good compromise between confocal imaging of thin cryosections (∼20 µm, e.g. (Wrobel *et al*, 2018; Huet *et al*, 2021) and light sheet microscopy of cleared intact cochleae (Keppeler *et al*, 2021). Indeed, the tick sections allow to preserve the anatomical integrity of the cochlea while providing a good antibody and light penetration for staining and imaging, respectively, two aspects that are critical for light sheet fluorescence microscopy.

Another critical aspect in the development of cell and gene therapy in the cochlea is the automated and reproducible evaluation of the effect of the therapy. In addition, preclinical development requires the evaluation of hundreds to thousands of samples. Therefore, we developed a pipeline in Arivis 4D using Cell Pose and a custom SGN model (Stringer *et al*, 2021; Pachitariu & Stringer, 2022) to segment and quantify SGN survival and transduction rate. This allowed a large number of samples to be processed using batch analysis with minimal time and human supervision.

### Design of the viral vector construct

First-in-human clinical data for cochlear gene therapy were recently reported for Otoferlin-related auditory synaptopathy patients who regained hearing by replacing the missing OTOF gene (Qi *et al*, 2024; Lv *et al*, 2024; Wang *et al*, 2024). These proof-of-concept studies help paving the way for the optogenetic hearing restoration.

Achieving efficient and safe ChR expression in adult SGNs is far from trivial and requires identification of the optimal combination of promoter, ChR, regulatory sequences, viral capsid, titer, and approach to viral vector delivery in the cochlea. Because the SGNs are not directly accessible, state-of-the-art molecular strategies (e.g., directed evolution strategies, (Jüttner *et al*, 2019; Cronin *et al*, 2014; Dalkara *et al*, 2013)) cannot be used to identify optimal viral constructs. Therefore, previous and current studies in the cochlea rely on in vivo screening of different viral constructs (Huet *et al*, 2021; Keppeler *et al*, 2018; Zerche *et al*, 2023; Bali *et al*, 2021). Lessons learned from previous works include: *i*) human synapsin (hSyn) promoter mediates efficient (Wrobel *et al*, 2018; Hernandez *et al*, 2014; Mager *et al*, 2018) and safe (Bali *et al*, 2022) ChR expression in the SGNs; *ii*) CatCh (Wrobel *et al*, 2018; Dieter *et al*, 2020; Keppeler *et al*, 2020; Michael *et al*, 2023) and f-

Chrimson (Mager *et al*, 2018; Bali *et al*, 2021) are two potential ChR candidates for hearing restoration; *iii*) ChR membrane expression can be restored after removing of GFP if replaced with trafficking sequences (Zerche *et al*, 2023); *iv*) f-Chrimson membrane expression is reduced when transduction occurs at adult age (Huet *et al*, 2021). In this study, hSyn was used in conjunction with CatCh fused to eYFP, for histological detection of CatCh in the tissue. To take into account the potential passage of viral vector suspension to the brain following cochlear administration (Talaei *et al*, 2019), that could lead to infection of central nervous system neurons, the capsid AAV.PHP-S was used for its narrow tropism towards the peripheral nervous system (Chan *et al*, 2017) and cochlear cells (Zhao *et al*, 2022). The viral vector suspension was administered at the titer at which it was produced: 6.4 x 10^12^ gene copies/mL, consistent with previous reports using adult gerbils (Huet *et al*, 2021; Wrobel *et al*, 2018; Dieter *et al*, 2019).

### Validating AAV-PHP.S to deliver ChR to the SGNs

The ability of AAV-PHP.S-hSyn-CatCh to drive ChR expression in the SGNs was first evaluated by intracochlear pressure injection in early postnatal age gerbils. AAV-PHP.S successfully drive ChR expression in the SGNs of both cochleae. Bilateral transduction following early postnatal injection is normal (Mager *et al*, 2018; Bali *et al*, 2021; Huet *et al*, 2021; Mittring *et al*, 2023; Richardson *et al*, 2021; Keppeler *et al*, 2018) and is discussed to occur through viral vector spread via the cochlear aqueduct and/or the endolymphatic duct to the cerebrospinal fluid space (Lalwani *et al*, 1996). Surprisingly, the SGN density was twofold decreased in the injected cochleae compared to the non-injected ones, suggesting a potential AAV dose- or protein level-dependent toxicity leading to SGN loss (Zerche *et al*, 2023). The precise folding and trafficking of proteins across membranes is complex but crucial, especially in the context of microbial opsin expression (Bedbrook *et al*, 2018). Impairment of these processes could potentially lead to toxic levels of cell stress through aggregation in the endoplasmic reticulum, trafficking defects and/or aggregate formation. Thus, cell-damaging stress may result from strong overexpression of the optogenetic actuator. Another surprise was to find IHC expressing GFP^+^ in the injected cochlea, even though IHCs do not natively express synapsin (Safieddine & Wenthold, 1999; Nouvian *et al*, 2011). Transcription of the protein in absence of a promoter specific to the IHCs is another suggestion of the high infection level occurring with AAV-PHP.S-hSyn-CatCh (6.4 x 10^12^ gene copies/mL). Optimization of the dose at which AAV-PHP.S-hSyn-CatCh should be injected, at a neonatal stage, to maximize transduction of SGNs was beyond the scope of this study. Nevertheless, these data suggest a relatively good tropism of AAV-PHP.S to capitalize on for gene delivery to adult SGNs.

### Improved viral vector-delivery to the adult SGNs using microcatheter round window administration and vent

Achieving reliable ChR expression in adult SGNs is key to clinical translation of the oCI. Previous work in adult gerbils found that pressure injection into the modiolus of AAV2/6-hSyn-CattCh enabled optogenetic modification of ∼30% of SGNs in approximately 40% of treated cochleae (Wrobel *et al*, 2018). Following work with other capsids and/or ChRs injected directly into the modiolus of gerbil cochleae, lower performance was reported in terms of transduction rate (∼10%) and proportion of successfully treated cochleae (∼10%, (Huet *et al*, 2021; Michael *et al*, 2023)). Alternatively, attempts to deliver the viral suspension directly into the perilymph either through the round window or via the posterior semicircular canal have not proven to be a reliable alternative (Richardson *et al*, 2021).

Is the direct injection of the AAV.PHP-S-hSyn-CatCh into the modiolus sufficient to improve the success of the treatment? Using it, oABRs could be measured in 20% of the treated cochleae for an overall transduction rate of 25.69% of all injected cochleae. Because of methodological differences, comparisons are complicated but it is reasonable to assume that AAV.PHP-S-hSyn-CatCh performs at least similarly than in previous reports (Wrobel *et al*, 2018; Michael *et al*, 2023). Nevertheless, this approach remains an interesting route, perhaps for cases where AAV re-dosing is required. It is worth noting that direct modiolus injection has been shown to be feasible in human temporal bone (Wrobel *et al*, 2021), where clear and visible landmarks are available.

As an alternative, we investigated if microcatheter pump-controlled administration approaches via the scala tympani could offer a viable preclinical alternative to the direct modiolus injection. Here, we demonstrated a fourfold increase in the success rate of measuring oABRs (80 vs 20), as well as an 8% increase in SGN survival (∼81 vs 73%) and a 1.75-fold increase in transduction rate (28 vs 16%) of the RW_µ-cat_ + vent compared to modiolus-injected cochleae. This improvement comes at the cost of a drastic loss of IHC, most likely due to the insertion and withdrawal of the catheter used to deliver the viral suspension (Jwair *et al*, 2023). The field of clinical cochlear implants has developed several robotic and automated approaches for minimally invasive insertion (De Seta *et al*, 2022). These approaches could be adapted for human catheter-based viral delivery if IHC survival is required. In a large fraction of treated cochleae with AAV.PHP-S-hSYn-CatCh, the remaining IHCs were expressing ChR. For both modiolus injection and microcatheter administration, the IHC transduction rate amounted to ∼38 and 30% for modiolus and RW_µ-cat_ + vent administration, respectively. The fact that no oABRs were measured from the modiolus-injected cochleae, where SGN transduction was close to 0%, supports that IHC optogenetic activation alone was not sufficient to drive synchronous firing in the SGNs. Moreover, the likelihood of viral particles reaching the SGNs via the IHCs seemed low, considering that pharmacologic depletion of the IHCs by intracochlear injection of kanamycin one week prior to viral vector delivery did not affect the performance of the procedure. Alternatively, co-transduction of IHCs and SGNs was found in RWµ-cat + vent treated cochleae, and their co-optical activation was associated with a slightly longer latency of the SGN compound action potential. As previously suggested, our data support that the contribution of optogenetically activated hair cells, if any, is very small compared to that of SGNs (Michael *et al*, 2023).

Why is scala tympani administration more efficient for transducing SGNs than direct injection into the modiolus where their cell bodies are located? One answer may be found in the available volume of the scala tympani: it is about 18 times larger than the modiolus (Keppeler *et al*, 2021), which is already densely packed with SGN cell bodies. Although the route by which the virus finds its way to the SGN can only be speculated at this point, our data suggest that an increased number of virus particles find their way to the SGN with RW_µ-cat_ + vent administration compared to direct modiolus injection. In the present study, a maximum volume of 5 µL was administered, corresponding to ∼80% of the cumulated volume of the scala timpani, vestibuli and media (Keppeler *et al*, 2021). This might suggest for mammals with bigger cochlea to scale up the volume of administered viral suspension to the volume of those cumulated cavities that would translate to ∼15 µL and ∼71 µL for guinea pig and humans, respectively.

### Lessons learned for optogenetic hearing restoration

This study indicates that AAV administration via the round window with pressure relief at the round window is a reliable approach to transduce SGNs. This approach has recently been successfully employed in clinical trials on AAV-mediated gene therapy of otoferlin-related auditory synaptopathy (Lv *et al*, 2024; Wang *et al*, 2024; Qi *et al*, 2024). There, IHCs have been the target that can be considered more accessible to virus administered to the *scala tympani* than SGNs that, except for their peripheral neurites innervating the organ of Corti, are thoroughly encased in the bony modiolus. Here we employed microcatheters that are smaller versions of catheters approved for clinical use and delivered volumes that relatively speaking come close to those applied to the human cochlea. Clearly, the reliability and efficacy of the optogene therapy need further optimization to provide optogenetic modification of the spiral ganglion in 100% of treated cochleae and in the majority of the SGNs. We note that 30% seems a good target of ChR^+^ positive cells given the experience of the more advanced vision restoration (Chaffiol *et al*, 2022). In fact, this expression rate has also supported demonstration of improved coding of spectral information in prior preclinical work on the waveguide or LED-based oCI stimulation of SGNs (Dieter *et al*, 2019, 2020). However, eventually, the benefits of oCI are predicted to be greatest with the largest possible population of ChR^+^ SGNs. Finally, our deafness model has limitations for it primarily represents acute deafness and might not resemble aspects of chronic deafness such as organ of Corti scarring and partial or complete loss of SGNs.

Off-target expression of the therapeutic construct, opsins in the case of optogenetic therapy, can cause adverse or confounding effects. We observed that despite choosing viral vector and promotor for specifically targeting SGNs, the only neural population with somatas contained in the cochlea, expression of ChR also occurred in hair cells to different extent. While this calls for further efforts for more selective optogenetic modification, it also provided us an opportunity to compare optogenetic stimulation of the auditory pathway with and without ChR expressing IHCs. This is relevant for future optogenetic hearing restoration in cases of residual low-frequency hearing currently treated with electroacoustic or hybrid stimulation of the eCI (Lenarz, 2017; Li *et al*, 2019). Our data suggest minimal impact of potential optogenetic stimulation of remaining IHCs on optically induced activity of the auditory pathway. We did not find differences in thresholds or amplitudes of the oABRs, but found a slightly delayed latency of the first oABR wave, which is consistent with optogenetically induced glutamate release from IHCs (Chakrabarti *et al*, 2022) contributing to SGN firing. Therefore, if remaining hair cells were transduced and optogenetically co-stimulated along with SGN, this might affect temporal coding and would need to be considered by sound coding strategies. The scope of our current study did not cover a comprehensive study of ChR expression across the brain or analysis of viscera. We note that different from a previous study employing early postnatal AAV injection into the mouse cochlea that showed ChR expression in various brain regions, the neural ChR expression upon microcatheter based AAV administration to the mature cochlea of the present study was largely limited to SGNs. This is promising and encourages comprehensive toxicology studies in preparation of clinical trials.

## Materials and methods

### Animals

Optogenetic data were obtained from 61 and control histological data from 22 Mongolian gerbils of either sex. For all procedures, animals were kept on a retro-controlled heating pad and their body temperature maintained at 37 degrees Celsius. All experiments were performed following the guidelines provided by the German national animal care and were approved by the board for animal welfare of the University Medical Center Göttingen and the animal welfare office of the state of Lower Saxony (LAVES). Animals were kept in a 12 hours light/dark cycle, with access to food and water *ad libitum*.

### Viral vector production

Viral vector employed in this study were produced as previously described (Huet & Rankovic, 2021). Briefly, pHelper plasmid (TaKaRa, USA), the trans-plasmid with PHP.S capsid or AAV2/9 capsid and the *cis*-plasmid with CatCh under the control of the human synapsin promotor were triple transfected with HEK-293T cells. Cells were regularly checked for mycoplasma contamination. Viral particles were collected 72 hours post-transfection from the medium and 120 hours post-transfection from both the cells and the medium. They were then treated by precipitation using 40% polyethylene glycol 8000 (Acros Organics, Germany) in 500 mM NaCl for 2 hours at 4°C. Following centrifugation, these particles were merged with cell pellets for subsequent processing. Cell pellets were suspended in 500 mM NaCl, 40 mM Tris, 2.5 mM MgCl2, pH 8, and 100 U/ml of salt-activated nuclease (Arcticzymes, USA) at 37 °C for 30 min. Subsequently, the cell lysates underwent clarification through centrifugation at 2000×g for 10 minutes and purification through iodixanol (Optiprep, Axis Shield, Norway) step gradients (15%, 25%, 40%, and 60%) at 350,000×g for 2.25 hours. The viruses were concentrated using Amicon filters (EMD, UFC910024) and then suspended in sterile phosphate-buffered saline (PBS) supplemented with 0.001% Pluronic F-68 (Gibco, Germany). Using an AAV titration kit (TaKaRa/Clontech), the viral vector titers were measured, by determining the number of DNase I resistant vg using qPCR (StepOne, Applied Biosystems). Silver stainings (Pierce, Germany) were used to check for the purity of the viruses routinely after gel electrophoresis (Novex™ 4–12% Tris–Glycine, Thermo Fisher Scientific). The presence of viral capsid proteins was then confirmed in all viral vector preparations. The viral stocks were stored at −80 °C until needed.

### Early postnatal injections in the *scala tympani*

Injections were performed as previously described (Huet & Rankovic, 2021; Huet *et al*, 2021). Briefly, under general Isoflurane anaesthesia (1.5 – 5%) and analgesia (subcutaneous injection of buprenorphine, 0.1 mg/kg and carprofen, 5 mg/kg), the left bulla of Mongolian gerbils of P7-8 was exposed via a retro-auricular incision. A volume of 1 – 1.5 µL of viral vector suspension, mixed with fast green (1:20), was loaded into a quartz micropipette (tip diameter approximately 20 µm, Science products; pulled with a P-2000 laser puller, Sutter Instruments) connected to a pressure microinjector (100-125 PSI, PLI-100 pico injector, Harvard Apparatus) and injected in the *scala tympani*. The right micropipette placement was confirmed visually by a perilymph reflux in the pipette tip when entering the *scala tympani*. After injection, the pipette was carefully retracted, the tissue above the injection site was repositioned and the wound was sutured. Animals were then daily tracked for the first 3 days and then weekly observed until oABRs were recorded. Carprofen (5 mg/kg) was given the first 3 days, and additional application could be performed at later point.

### Adult modiolus injections

Injections were performed as first described by Wrobel and colleagues (Wrobel *et al*, 2018). Briefly, under general Isoflurane anaesthesia (1.5 – 5%) and analgesia (subcutaneous injection of buprenorphine, 0.1 mg/kg and carprofen, 5 mg/kg), the left bulla of Mongolian gerbils of at least 8 weeks was exposed via a retro-auricular incision. A bullostomy was performed to expose the round window niche and a hole was drilled in the modiolus. A volume of 3 µL of viral vector suspension was subsequentially injected in the drilled hole using a quartz micropipette (same design as described in early postnatal injection). After the injection, the tissue above the bulla was repositioned and the wound was sutured. Animals were then daily tracked for the first 3 days and received additional subdermal carprofen (5 mg/kg) injections. Later, animals were weekly tracked.

### Adult *scala tympani* administrations

Administrations were performed similarly than the modiolus injection previously described at the difference that no hole was drilled in the modiolus. In this case, a microcatheter provided by the cochlear implant manufacturer MED-EL was filled with 5 µL of viral vector suspension filled with fast green (1:20) and connected to a micro-infusion pump (UltraMicroPump3, Word Precision Instrument, United States of America). If required by the procedure, an evacuation vent was drilled next to the oval window or in the posterior semi-circular canal. After insertion of the microcatheter in the RW (4.5 – 5.5 µm deep), the viral vector was administered at 250 – 300 nL/min and the administration was stopped either following reflux of the viral suspension or when the 5 µL were dispensed. Typically, the administration took 20 mins. Following careful retraction of the microcatheter, the procedure was finished as described for the modiolus injections.

### Deafening

The deafening procedure was achieved under general Isoflurane anaesthesia (1.5 – 5%) and analgesia (subcutaneous injection of buprenorphine, 0.1 mg/kg and carprofen, 5 mg/kg). The left bulla of Mongolian gerbils of at least 8 weeks was exposed via a retro-auricular incision and a bullostomy was performed to expose the round window niche. A volume of 3 µL of kanamycin (100 mg/ml, CarlRoth Gmbh, T832.2) was subsequentially injected through the round window membrane using a quartz micropipette (same design as described in early postnatal and adult modiolus injection). After the injection, the tissue above the bulla was repositioned and the wound was sutured. Animals were then daily tracked for the first 3 days and received additional subdermal carprofen (5 mg/kg) injections. From the second week after injection, animals were weekly tracked and additional application of carprofen (5 mg/kg) could be performed.

### Stimulation

Stimuli were generated via a custom-made system based on NI-DAQ-Cards (NI PCI-6229, National Instrument, Austin, USA) and custom-written MATLAB scripts (The MathWorks, Natick, USA). Acoustic stimuli were generated at a sampling rate of 830 kHz and presented via an open-field speaker (Avisoft Inc., Germany) placed at ∼15 cm from the left pinna. A ¼ inch microphone and amplifier (D4039; 2610; Brüel & Kjaer GmbH, Naerum, Denmark) were used to calibrate sounds. Optical stimuli were generated at a sampling rate of 50 kHz and were delivered into the cochle via an optical fiber (ø = 200 µm, 0.39 NA, Thorlabs GmbH, Germany) coupled to a blue laser (473 nm, MLLFN-473-100, 100 mW, Changchun New Industry Optoelectronics). The maximum radiant flux at the output of the optical fiber was calibrated before every experiment (LaserCheck; Coherent Inc.) and later used for calibration. Access to the round window was gained similarly than for the viral vector injection described above. Stimulations were presented at 20 Hz at least 500 times per tested condition for auditory brainstem stimulation and 100 times for compound action potentials.

### Auditory brainstem recordings

At least 8 weeks after early postnatal injection and 4 weeks after adult injection/administration, auditory brainstem responses (ABR) were recorded. The ABRs were recorded using a custom-made differential amplifier with sub-dermal needle electrodes placed at the vertex, below the ipsilateral pinna and at the contralateral leg. Potentials were digitalized using the same NI-DAQ-Cards than described above for stimulation at a sampling rate of 50 kHz. Response to every stimulation presentation were saved for offline analysis. ABR analysis was performed using custom-made Matlab scripts: traces were filtered (2^nd^ order Butterworth pass-band filter between 300 and 3000 Hz) and averaged per tested condition. Wave detection (wave I, II and III) was achieved manually and the activation threshold defined as the condition for which the smallest ABR could be visually detected. Amplitude of the different waves was obtained by detecting automatically the negative peak following each wave.

### Compound action potential recordings

For a subset of animals, compound action potentials (CAP) were recorded using a custom-made differential amplifier with a silver ball electrode place at the round window niche and two sub-dermal needle electrodes placed below the ipsilateral pinna and at the contralateral leg. Potentials were digitalized using the same NI-DAQ-Cards than described above for stimulation at a sampling rate of 50 kHz. Response to every stimulation presentation were saved for offline analysis. ABR analysis was performed using custom-made Matlab scripts: traces were filtered (2^nd^ order Butterworth pass-band filter between 100 and 10000 Hz) and averaged per tested condition. The first negative was automatically detected if its amplitude was lower than −1 µV and the activation threshold defined as the condition for which the smallest CAP was detected.

### Cochlea and brain harvesting

At the end of the CAP/ABR recordings, animals were sacrificed by cervical dislocation under deep anaesthesia and their cochlea and brain were immediately collected and fixed in formaldehyde 3.7% (Carl Roth, Germany). Tissue fixation was done for 1 hour and at least for 24 hours for cochleae and brains, respectively. Cochleae were then decalcified in 0.12 M EDTA for 7 days.

### Mid-modiolar cryosection of the cochlea

Cochleae were first dehydrated by incubating them overnight in 30% sucrose solution in PBS. Next, slices were embedded in Cryomatrix (Epredia, United Stated of America) and 25 µm slices were obtained using a cryostat (Leica CM3050S, Germany) using a cutting angle parallel to the modiolus. After staining, slices were mounted with Mowiol 4-88 (Carl Roth, Germany) and imaged using a LSM 510 Zeiss confocal microscope (Zeiss, Germany) with a 40x objective. A z-stack image (1 µm z-resolution) was taken per cochlear turn.

### Cross-modiolar section of the cochlea

Cochleae were sliced using a vibratome (Leica 1200S speed = 0.02 mm/s, amplitude = 0.6 mm) with a thickness of 220 µm at a cutting angle perpendicular to the modiolus. For staining, one slice per cochlear turn was kept on the basis of visualizing the IHCs and the SGNs innervating them. After staining, slices were cleared and mounted in FOCM (Zhu *et al*, 2019). A 30 µm z-stack (SGNs: 1 µm z-resolution; IHCs: 3 µm z-resolution) from the SGNs and IHCs were taken using a Leica SP8 confocal microscope (Leica microsystems, Germany) with a 40x objective. Overview images were taken using the same microscope equipped with a 20x objective and using the inbuilt tile function.

### Cochlear histology

Permeabilization and blocking of the slices was obtained by 2 hours incubation in 16% goat serum dilution buffer (16% normal goat serum, 450 mM NaCl, 0.6% Triton X-100 20 mM phosphate buffer, pH 7.4). Information about the primary antibody incubation is summarized in table 3. Staining with secondary antibodies was done in one hour for mid-modiolar cryosections and 48 hours for cross-modiolar sections at 4°C using goat anti-chicken 488 (A11039, Life Technologies, 1:200), goat anti-guinea-pig 568 (A11075, Life Technologies, 1:200) and goat anti-rabbit 647 (A21244, Life Technologies, 1:200).

**Table 3.** Summary of primary antibody incubation per slicing approach.

| Marker of | Primary antibody reference | Mid-modiolar cryosection | Cross-modiolar section |
| --- | --- | --- | --- |
| GFP | chicken anti-GFP #ab13970, Abcam, USA | 1:500, overnight at 4°C | 1:500, 72 hours at 4°C |
| SGNs | guinea pig anti-parvalbumin 195004, Synaptic Systems, Germany | 1:300, overnight at 4°C | 1:200, 72 hours at 4°C |
| IHCs | rabbit anti-calretinin 7697, Swant, Switzerland | NA | 1:500, 72 hours at 4°C |

### Histological analysis

Analysis of mid-modiolar images was done using a custom-made script in Matlab as previously described in (Mittring *et al*, 2023; Huet *et al*, 2021). The conversion from SGN density in 2D to 3D was done by multiplying the surface in which SGN were segmented by the number of stack that were imaged and the z-resolution.

Analysis of cross-modiolar images was done within Vision 4D (Zeiss, Germany). SGN somas were automatically segmented using a custom model in Cellpose 2.0 (Pachitariu & Stringer, 2022; Stringer *et al*, 2021). The volumetric boundary containing all segmented SGNs was then computed using a custom-written function. Identification of the GFP positive SGNs was done as previously described in (Mittring *et al*, 2023; Huet *et al*, 2021), using a python script directly in Vision 4D. IHCs and ChR-expressing IHCs were counted manually in Fiji from the maximum projection along a manually drawn line. ChR-expressing IHCs were visually identified as cells brighter than the mean + 2 standard deviation of the GFP background signal measured from a non-neural region of the image.

### Data analysis and statistics

All data were analyzed using MATLAB (MathWorks). Averages were expressed in figures and main text as mean ± SEM. For statistical comparison between two independent groups, data were tested for normality (Jarque-Bera test) and Student’s t-test or Wilcoxon rank sum test were used accordingly. For statistical comparison between more than two independent groups, data were tested for normality (Jarque-Bera test), and one-way ANOVA or a Kruskal-Wallis test was applied accordingly following by a Tukey-Kramer post-hoc test.

## Author contribution

Anupriya Thirumalai: Conceptualization, Validation, Formal analysis, Investigation, Data curation, Writing review & editing, Visualization.

Jana Henseler: Investigation Marzieh Enayati: Investigation

Kathrin Kusch: Viral vector design and production. Roland Hessler: Microcatheter design and production.

Tobias Moser: Conceptualization, Validation, Resources, writing original draft, Writing e review & editing, Supervision, Project administration, Funding acquisition.

Antoine Tarquin Huet: Conceptualization, Validation, Formal analysis, Investigation, Resources, Data curation, writing original draft, writing review & editing, Supervision, Project administration, Funding acquisition.

## Acknowledgement

We thank Daniela Gerke for supporting us with the viral vector production and in performing the immunolabeled mid-modiolar cochlear cryosection preparation. We thank Christiane Senger-Freitag for helping us in performing viral vector administration on to the gerbils at the postnatal stage, Gerhard Hoch for the engineering support and Patricia Räke-Kügler for excellent administrative support. Additionally, we are also grateful for Elena Kleemann, Thomas Dubois and Edouard Toussaint for giving us extra support with the immunolabelling of the brain slices and for some of the cochlear samples taken as the control using the cross-modioloar sectioning. Additionally, we are very grateful to Dr. Bettina Wolf for her involvement in this study, especially at the initial stages with critical decision making, and assistance. We would like to thank the MED-EL company and Dr. Roland Hessler for providing us with the micro catheters. We would like to acknowledge funding by the DFG priority programmme SPP 1926 - “Next Generation Optogenetics”, the Cluster of Excellence (EXC2067) Multiscale Bioimaging (AH and TM) as well as by the European Research Council through the Advanced Grant “DynaHear” under the European Union’s Horizon 2020 Research and Innovation program (grant agreement No. 101054467, TM) and by Fondation Pour l’Audition (FPA RD-2020-10 TM). Lastly, this work was conducted as part of the PhD project of A. Thirumalai at the University of Göttingen (GGNB). Hence, the authors thank the University of Göttingen for their support.

---

**Figure S1.**
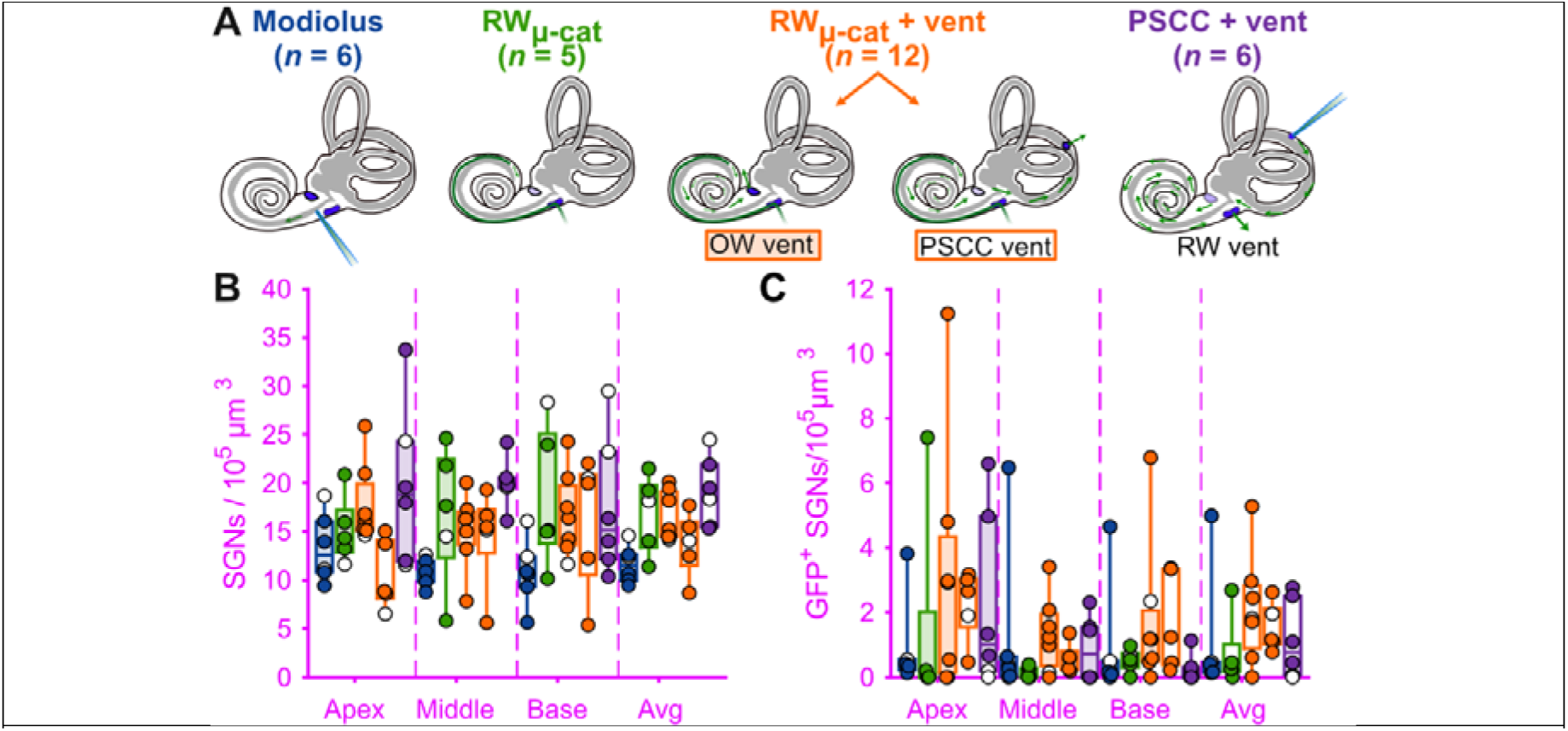
Comparison of viral administration approaches in the adult gerbil cochlea. **A.** Schematic representation of the different administration approaches (see Materials and methods for details). **B-C.** Quantification of the SGN density (B) and GFP^+^ SGN density (C) from the injected cochleae with the different administration approaches presented in A. Filled makers were used when positive oABRs were measured and an open-marker for the negative oABRs. Box plots show minimum, 25^th^ percentile, median, 75^th^ percentile, and maximum. Averaged ± SEM. Kruskal-Wallis test followed by Tukey-Kramer post-hoc test (*, *P* ≤ 0.05; **, *P* ≤ 0.01).

**Figure S2.**
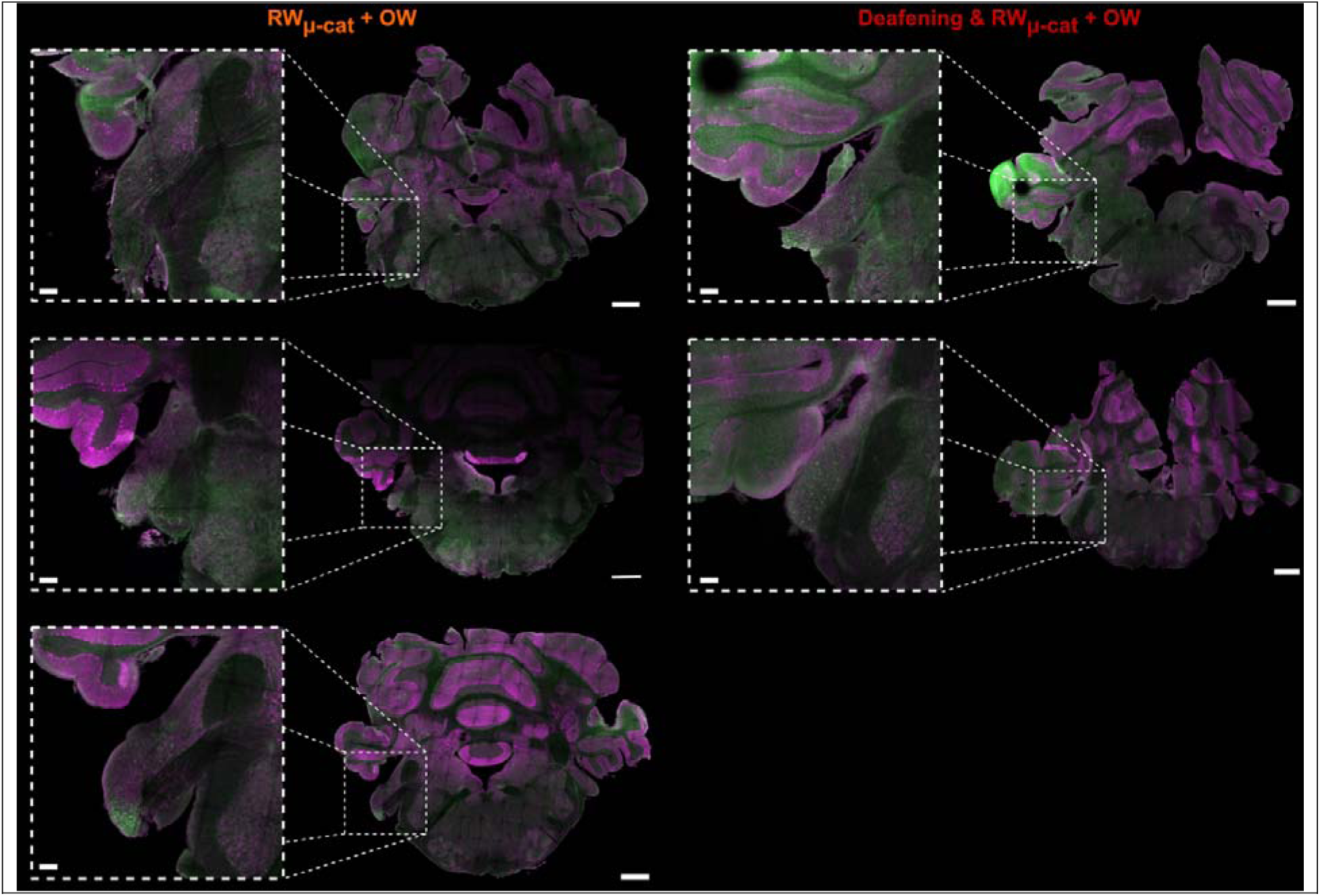
Absence of transduction in the central nervous system following RW_µ-cat_ + vent AAV-administration. Coronal slices of 5 gerbil brains following RW_µ-cat_ + vent AAV-administration. Slices were stained for parvalbumin (purple) and GFP (green), scale bar = 1 mm. The insert shows a magnification of the antero-ventral cochlear nucleus where GFP signal is found in axons of the SGNs but not in the neurons on which they project (scale bar = 0.2 mm).

